# The kynurenine pathway is essential for rhodoquinone biosynthesis in *Caenorhabditis elegans*

**DOI:** 10.1101/645929

**Authors:** Paloma M. Roberts Buceta, Laura Romanelli-Cedrez, Shannon J. Babcock, Helen Xun, Miranda L. VonPaige, Thomas W. Higley, Tyler D. Schlatter, Dakota C. Davis, Julia A. Drexelius, John C. Culver, Inés Carrera, Jennifer N. Shepherd, Gustavo Salinas

**Author notes:** To whom correspondence should be addressed: Jennifer N. Shepherd: Department of Chemistry and Biochemistry, Gonzaga University, 502 E. Boone Ave., Spokane, WA 99258, Gustavo Salinas: ^2^Laboratorio de Biología de Gusanos, Institut Pasteur de Montevideo, Uruguay, Mataojo 2020, Montevideo, Uruguay 11400. Oregon Health Sciences University, 3181 SW Sam Jackson Park Rd., Portland, OR 97239. Johns Hopkins University School of Medicine, 733 N. Broadway, Baltimore, MD 21205. Multicare Allenmore Hospital, 1901 S. Union Ave., Tacoma, WA 98405. Departamento de Ciencias Farmacéuticas, Área Farmacología, Facultad de Química, Universidad de la República, Montevideo, Uruguay 11800 (Inés Carrera).

## Abstract

A key metabolic adaptation for some species that face hypoxia as part of their life-cycle involves an alternative electron transport chain in which rhodoquinone (RQ) is required for fumarate reduction and ATP production. RQ biosynthesis in bacteria and protists requires ubiquinone (Q) as a precursor. In contrast, Q is not a precursor for RQ biosynthesis in animals such as parasitic helminths, and this pathway has remained elusive. We used *Caenorhabditis elegans* as a model animal to elucidate several key steps in RQ biosynthesis. Through RNA interference and a series of mutants, we found that arylamine metabolites from the kynurenine pathway are essential precursors for RQ biosynthesis *de novo*. Deletion of *kynu-1*, which encodes a kynureninase that converts L-kynurenine (KYN) into anthranilic acid (AA), and 3-hydroxykynurenine (HKYN) into 3-hydroxyanthranilic acid (3HAA), completely abolishes RQ biosynthesis, but does not affect Q levels. Deletion of *kmo-1*, which encodes a kynurenine 3-monooxygenase that converts KYN to HKYN, drastically reduces RQ, but not Q levels. Knockdown of the Q biosynthetic genes, *coq-5* and *coq-6*, affects both Q and RQ levels demonstrating that common enzymes are used in both biosynthetic pathways. Our study reveals that two pathways for RQ biosynthesis have independently evolved. In contrast to bacteria, where amination is the last step in RQ biosynthesis, worms begin with the arylamine precursor, AA or 3HAA. Since RQ is absent in mammalian hosts of helminths, inhibition of RQ biosynthesis may have broad implications for targeting parasitic infections which cause important neglected tropical diseases.

---

Adaptation to hypoxia is crucial to survival in several animal lineages (1). Such is the case with helminths (parasitic nematodes and platyhelminths), which are facultative anaerobes, and live part of their life-cycle under low oxygen tension in the gastrointestinal tract of their vertebrate hosts. One of the key adaptations of these lineages is the use of an alternative electron transport chain (ETC) that allows them to harvest energy under hypoxic conditions (2-4). In the absence of oxygen, complex II of this alternative ETC functions in the opposite direction to the conventional ETC. To facilitate this reversal, fumarate is used as the final electron acceptor, and rhodoquinone (RQ) functions as the electron transporter. RQ differs from ubiquinone (Q), the electron transporter in the conventional ETC, by one substituent on the benzoquinone ring: RQ has a 2-amino substituent, while Q has a methoxy group in this position (Fig. 1A). RQ has a lower redox potential than Q (−63 mV versus 110 mV, respectively) (5, 6), enabling RQ to receive electrons from NADH through complex I and donate them to fumarate though complex II (Fig. 1B) (3, 7). In contrast to other fermentative pathways, the alternative ETC allows proton pumping and ATP synthesis through complex V, leading to higher efficiency in harvesting energy. This pathway, in which RQ is the signature metabolite, is also found in some bacteria and protists (1).

**Figure 1.**
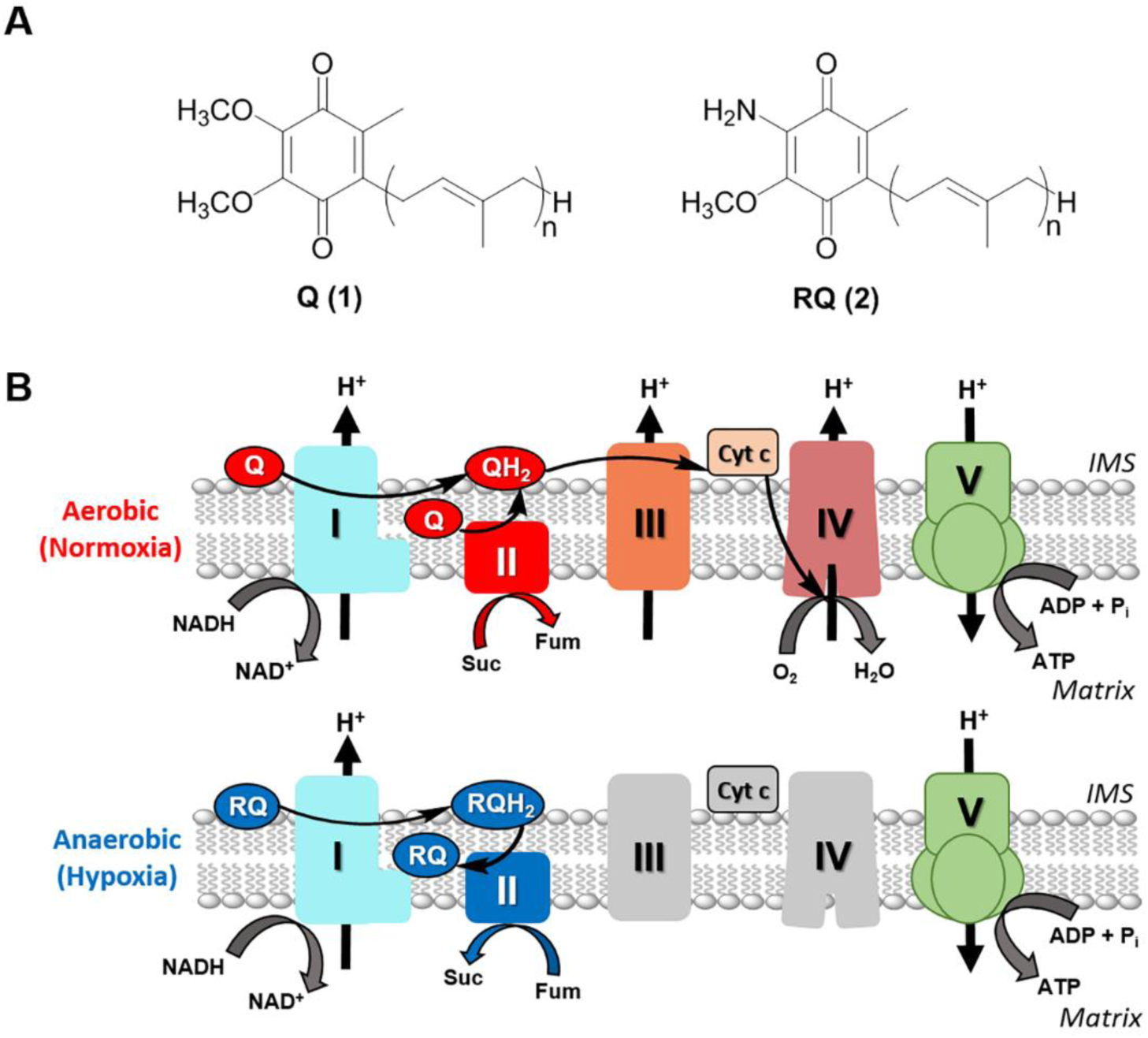
Structure and function of rhodoquinone (RQ) and ubiquinone (Q) in the mitochondria of helminths. (A) Structures of ubiquinone (UQ or Q, compound 1) and RQ (compound 2), where n varies from 6-10 depending on species. (B) Q and RQ are part of the mitochondrial electron transport chains (ETC) in normoxia and hypoxia, respectively. In normoxia, electrons from NADH and succinate are transferred to Q through complexes I and II, then from QH_2_ to cytochrome c (cyt c) through complex III, and finally from cyt c to O_2_ through complex IV. In hypoxia the ETC functions with complexes I and II only. Electrons are transferred from NADH to RQ through complex I and then from RQH_2_ to fumarate through complex II. In both ETCs, a proton gradient across the inner membrane is generated and used to produce ATP through complex V.

The biosynthetic pathway of RQ has been extensively studied in *Rhodospirillum rubrum*. In this organism, RQ biosynthesis requires Q as a precursor (8). Subsequently, it was discovered that the *R. rubrum rquA* gene is essential for RQ, but not for Q biosynthesis (9). Despite a comprehensive study of *R. rubrum* knock-out mutants, using aerobic versus anaerobic transcriptome data and comparative genomic analysis, no other genes besides *rquA* were identified to be essential for RQ biosynthesis (10). In parallel, it was shown that unicellular eukaryotes also possess a homolog of the *rquA* gene, most likely acquired by horizontal gene transfer (11). These studies indicate that *rquA* is the gene signature for RQ biosynthesis in bacteria and protists. More recently, the heterologous expression of *R. rubrum rquA* in two non-RQ-producing species, *Escherichia coli* and *Saccharomyces cerevisiae*, resulted in the *in vivo* conversion of native Q to synthetic RQ (12). Despite these advances, the biosynthesis of RQ in animals has not been elucidated, and the key genes involved have remained elusive.

RQ has been found in all helminths where it has been examined (7, 13). Importantly, RQ is also synthesized by the free-living nematode *Caenorhabditis elegans* (14), which faces hypoxia during development or as an environmental challenge. Studies in *C. elegans* have shown that Q is not a required precursor of RQ. Indeed, a KO strain in *coq-7* (also known as *clk-1*) abolishes Q biosynthesis without affecting RQ biosynthesis (15, 16). While helminths are not easily approachable, *C. elegans* is a formidable experimental organism (17), and has been used as a model for parasitic nematodes (18). In this study, we elucidate key steps in the RQ biosynthesis pathway using *C. elegans*. We demonstrate that the kynureninase KYNU-1 is essential for RQ biosynthesis, and based on RNAi experiments, we propose that RQ and Q have parallel pathways starting from different precursors. Since RQ is not synthesized or used by mammalian hosts, but required for parasite survival, the RQ biosynthetic pathway is a unique target for antihelminthic drug design.

## Results

### *KYNU-1 is essential for RQ biosynthesis in* C. elegans

Due to the differences in RQ biosynthesis in *R. rubrum* and *C. elegans*, we reasoned that, in the case of animals, the amino group at position 2 of the benzoquinone ring (Fig. 1A), may be added at the beginning of RQ biosynthesis, rather than at the end. Since *kynu-1* encodes a kynureninase that catalyzes the synthesis of two arylamines, anthranilic acid (AA) and 3-hydroxyanthranilic acid (3HAA), from *L*-kynurenine (KYN) and 3-hydroxy *L*-kynurenine (3HKYN), respectively (Fig. 2A), we examined RQ biosynthesis in a *kynu-1* KO strain. No trace of RQ was observed in the KO animals (Fig. 2B). In contrast, Q levels were not reduced in the KO. RNAi-mediated knockdown of *kynu-1* in the *C. elegans* strain *rrf-3*(pk1426), which is hypersensitive to RNAi (19), exhibited a significant decrease in RQ levels (p < 0.001), with no decrease in Q, as compared to the empty vector (EV) control (Fig. 2B). The expression of the wild-type *kynu-1* allele in the *kynu-1* strain under the control of its own promoter rescued RQ biosynthesis (Fig. 2B). These results allow us to conclusively demonstrate that *kynu-1* is essential for RQ biosynthesis, and strongly suggest that AA or 3HAA are key precursors for RQ biosynthesis. Intriguingly, supplementation experiments with AA or 3HAA did not restore RQ biosynthesis in the *kynu-1* strain (Fig. 2B). These results indicate that RQ is synthesized *de novo* in *C. elegans*.

**Figure 2.**
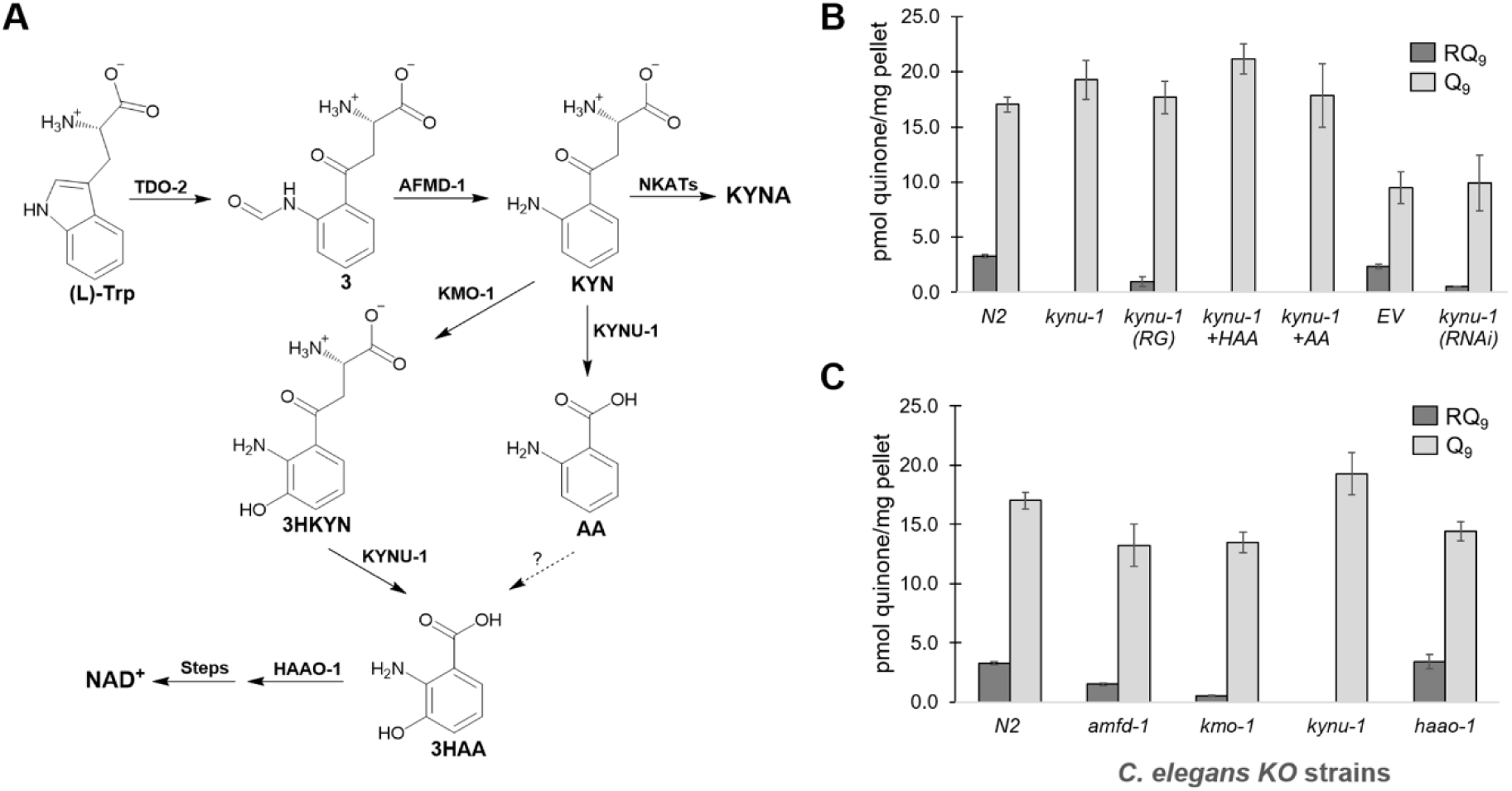
The kynurenine pathway is essential for RQ biosynthesis. (A) In the kynurenine pathway, *L*-tryptophan is first converted to *L*-formyl kynurenine (compound 3) by tryptophan 2,3-dioxygenase (TDO-2), which is then converted to kynurenine (KYN) by an arylformamidase (AFMD-1). Kynurenine (KYN) is a branch point and can be converted to 1) kynurenic acid (KYNA) by kynurenine aminotransferases (NKATs); 2) 3-hydroxykynurenine (3HKYN) by kynurenine 3-monooxygenase (KMO-1); or 3) anthranilic acid (AA) by the kynureninase (KYNU-1). KYNU-1 also transforms 3HKYN to 3-hydroxyanthranilic acid (3HAA), which has also been proposed to form from AA. Finally, 3HAA is converted to 2-amino-3-carboxymuconic semialdehyde (not shown) by 3-hydroxy anthranilic acid oxygenase (HAAO-1), which is then converted to NAD^+^. (B) Deletion of *kynu-1* from N2 *C. elegans* abolished RQ biosynthesis. Overexpression of *kynu-1* wild-type allele in the mutant *kynu-1* strain restored RQ biosynthesis (RG). Supplementation with 3HAA and AA did not rescue RQ levels. The *kynu-1* RNAi significantly reduced RQ levels in comparison to the empty vector (EV) control in *rrf-3*(pk1426) worms. (C) *amfd-1* and *kymo-1* strains significantly reduced RQ levels, compared to N2, while the *haao-1* strain had no effect. A full statistical analysis is given in Table S1.

Since KYN can be converted to 3HKYN by kynurenine 3-monooxygenase, KMO-1, we analyzed a *kmo-1* mutant strain in order to discriminate whether AA or 3HAA is the RQ precursor. The *kmo-1* strain had significantly reduced RQ levels compared to N2 (p < 0.001), but RQ biosynthesis was not completely abolished (Fig. 2C). The result is consistent with the fact that a hydroxyl substituent can be introduced at position 3 of the aromatic ring by *kmo-1*-dependent and *kmo-1*-independent routes (20, 21). Since KYNU-1 is required for the biosynthesis of both metabolites, this would explain the absolute requirement of this gene. Since both *kynu-1* and *kmo-1* belong to the kynurenine pathway (Fig 2A), we analyzed a KO mutant in *afmd-1*, the gene which precedes *kynu-1* in the pathway. *afmd-1* encodes the kynurenine formamidase that converts *L*-formyl kynurenine to KYN. Surprisingly, the *afmd-1* strain only reduced RQ levels to about half that of N2 (p < 0.001). We also analyzed a mutant strain in *haao-1*, which encodes a 3-hydroxyanthranilic acid oxygenase (HAAO-1) downstream of KYNU-1 in the pathway (Fig. 2A). This strain did not affect RQ biosynthesis (Fig. 2C). Thus, 2-amino-3-carboxymuconic semialdehyde is unlikely to be a precursor for RQ biosynthesis. Instead, RQ biosynthesis most likely branches from the AA or 3HAA precursors in the kynurenine pathway.

### kynu-1 *expression is restricted to the hypodermis of the worm*

To assess the expression pattern of *kynu-1* during *C. elegans* lifecycle, we generated and analyzed a translational reporter strain expressing GFP under the control of the *kynu-1* promoter (IH25 strain). We detected expression of *kynu-1* in the embryo early in the E lineage and epidermis. This pattern was maintained in the L1 stage but starting at L2 through adulthood expression was only seen in the epidermis (Fig. 3 and Fig. S1). Since RQ is supposed to play a key role as part of an alternative ETC under hypoxia, we analyzed whether expression of *kynu-1* was affected after exposure to 0.4% oxygen during 24 h in adult worms. We did not observe any obvious increase in GFP fluorescence nor a difference in the spatial expression of the reporter. This suggests that *kynu-1* expression is not regulated under hypoxic conditions.

**Figure 3.**
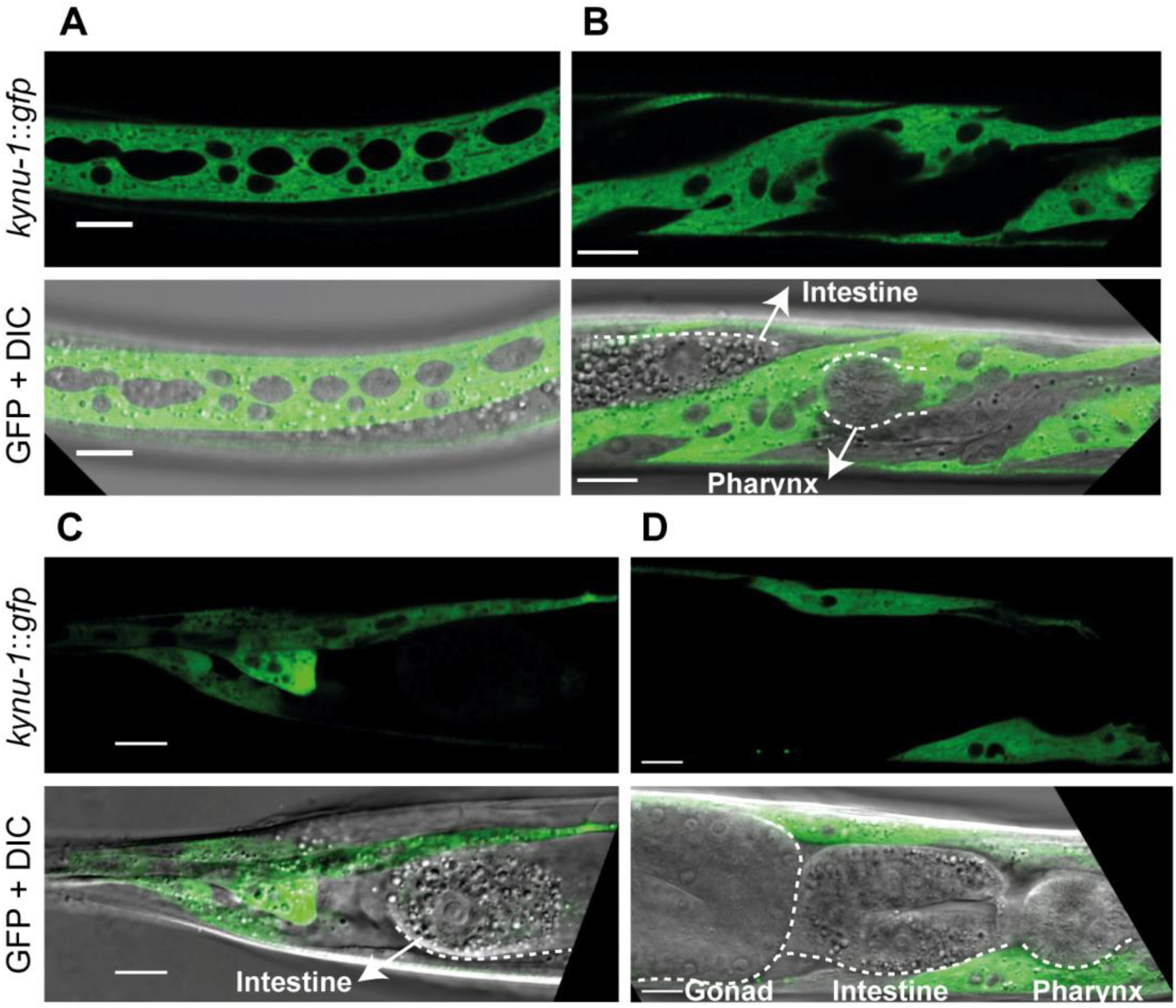
*kynu-1* is expressed in the hypodermis with cytosolic localization. Confocal images of selected planes show head, middle body and tail regions of P*kynu-1*::*kynu-1*::*gfp* transgenic animal expression. (A) Middle body region, (B) Head of an L3 worm (lateral views), (C) Head and (D) Tail of an adult worm (lateral views). Pharynx, intestine and gonad are indicated. Scale bar: 10 μm.

### Enzymes involved in Q biosynthesis are also involved in RQ biosynthesis

In the case of *S. cerevisiae*, Q can be synthesized from 4-hydroxybenzoic acid (4HB) or 4-aminobenzoic acid (pABA) in parallel pathways using common enzymes in most steps (22, 23) (Fig. 4A). Thus, we reasoned that some of the enzymes may also be involved in RQ biosynthesis. We performed RNAi knockdown assays of *coq-3, coq-5, coq-6* and *coq-7* genes. We found that *coq-5* and *coq-6* RNAi significantly decreased both Q and RQ levels compared to controls (p < 0.001), while *coq-3* had a smaller effect on both (Fig. 4B and Table S1). As expected, *coq-7* RNAi significantly decreased Q levels (p = 0.010), but not RQ. The mRNA levels for the silenced genes indicated efficient interference in all but the *coq-3* RNAi samples (Fig. S2). These results clearly indicate that COQ-5 and COQ-6 are involved in both Q and RQ biosynthesis. Our results support the existence of parallel pathways that use several common enzymes to synthesize Q and RQ from different precursors.

**Figure 4.**
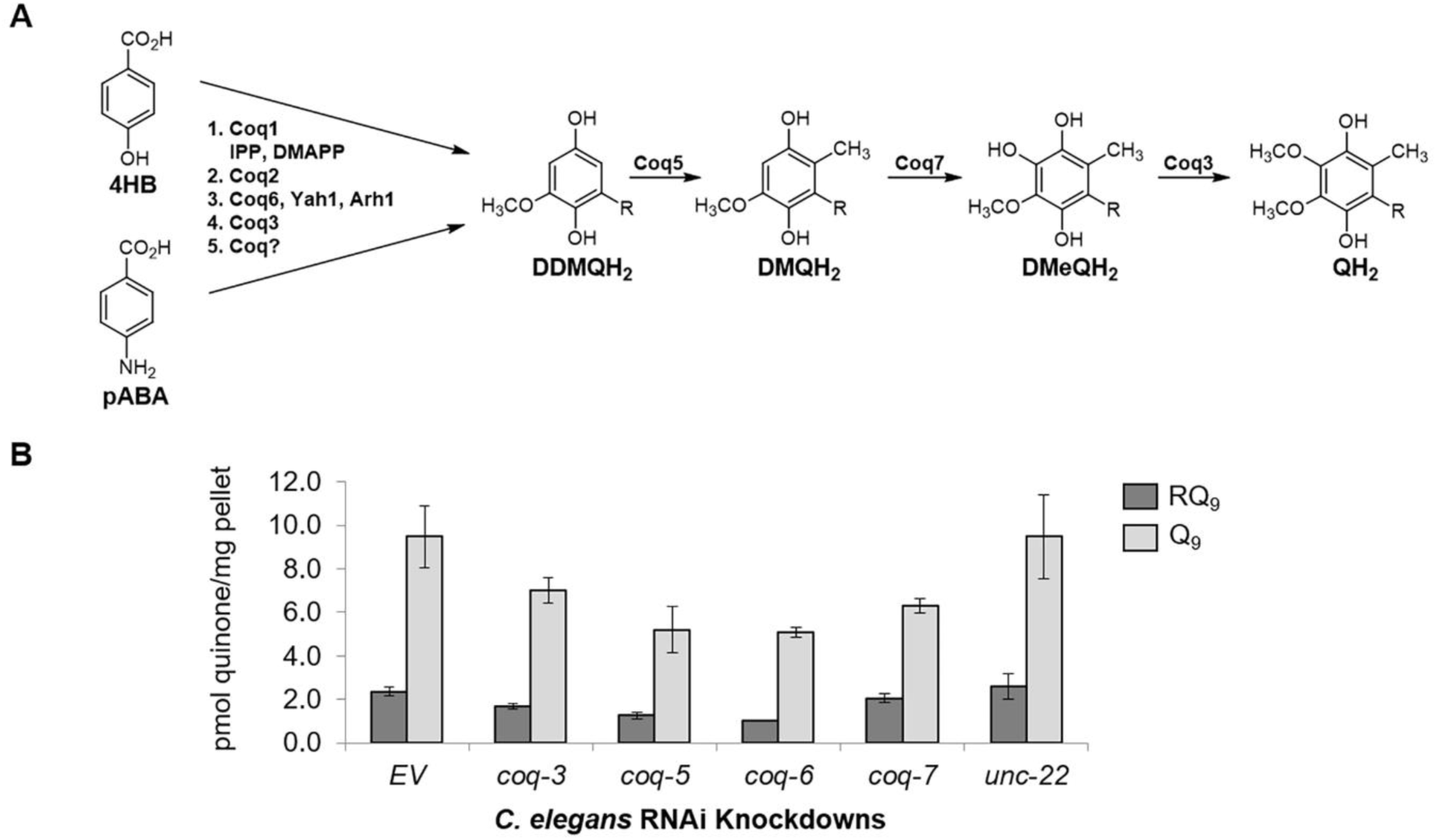
The biosynthesis of RQ shares common enzymes with the Q biosynthetic pathway. (A) The Q biosynthetic pathway in yeast can start from either 4-hydroxybenzoic acid (4HB) or *para*-aminobenzoic acid (pABA). These pathways share the enzymes Coq1, Coq2, Coq6 (with Yah1 and Arh1), and Coq3. They merge at the common precursor demethyldemethoxyubiquinone (DDMQH_2_), which is converted to QH_2_ in three steps by Coq5, Coq7 and Coq3, respectively. (B) RNAi strains of *coq-3, coq-5*, and *coq-6 C. elegans* show significant reduction of both RQ and Q, as compared to the empty vector (EV) and *unc-22* controls (Table S1). RNAi of *coq-7* significantly reduces Q levels, but RQ biosynthesis is unaffected (Table S1).

## Discussion

The biosynthesis of RQ in animals has remained a puzzle for decades (24). In bacteria and protists, RQ is derived from Q, and *rquA* is the gene signature for its biosynthesis (9, 11). In contrast, in animals, RQ is not derived from Q, and no RQ-specific gene has been discovered. By analogy to the biosynthetic pathway of Q in yeast from pABA (22), we reasoned that the 2-amino substituent of RQ could be derived from an arylamine precursor. While this manuscript was in preparation, a different group independently reported the essential role of KYNU-1 for RQ biosynthesis, using the *kynu-1* strain CB1003 (25). Our study was performed with the *kynu-1* strain Tm4924. The genetic rescue of Tm4924 and RNAi experiments that we performed confirmed this finding. These results indicate that AA and/or 3HAA are RQ precursors. Consistent with this view, the strain used in this study has been previously reported to show increased levels of KYN and 3HKYN (21). Interestingly, supplementation with AA or 3HAA did not rescue RQ biosynthesis, suggesting the absence of transporters for uptake of these metabolites. The *kmo-1* strain showed significantly reduced levels of RQ. Thus, whether AA or 3HAA or both are precursors of RQ is unclear. The *kynu-1-*dependent, *kmo-1*-independent biosynthesis of 3HAA has been postulated in several studies (20, 21), but to the best of our knowledge, no clear evidence regarding this reaction has been reported. In any case, the drastic decrease of RQ biosynthesis in *kmo-1* strain would suggest that 3HAA is a RQ precursor. The mutant strain in *afmd-1*, upstream of *kynu-1*, did not completely abolish RQ biosynthesis. This result would be explained if the *afmd-1* strain is not a null-mutant or if KYN can be acquired from *E. coli*

KYNU-1 expression was mostly restricted to the hypodermis, suggesting that the precursor of RQ is transported to other tissues. Two genes of the kynurenine pathway, *tdo-2* and *kmo-1*, have also been found to be expressed almost exclusively in the worm hypodermis (26). KYN, AA and 3HAA transport to other worm tissues is likely to be highly relevant since they are also precursors for other key metabolites, such as quinolinic acid and kynurenic acid. Interestingly, enigmatic deposits of fluorescent AA glycosyl esters are found in gut granules in dying worms (20). We found that the expression of *kynu-1* was not upregulated under hypoxic conditions. KYNU-1 may be constitutively expressed since it is also essential for *de novo* synthesis of NAD^+^ (27). An important conclusion of our study is that the kynurenine pathway is a complex metabolic hub, and that AA or 3HAA is a likely branch point for RQ biosynthesis.

Our study reveals that from the kynurenine pathway branch point, Q and RQ biosynthesis in *C. elegans* make use of common enzymes, since *coq-5* and *coq-6* RNAi led to a significant decrease of both quinones. The fact that the enzymes involved in Q biosynthesis do not have strict substrate specificity is highlighted by the parallel pathways of Q biosynthesis in yeast that start from different precursors (23). In addition, neo-functionalization of Coq enzymes has been described for the COQ-5 bacterial ortholog (UbiE/MenG) in the biosynthesis of menaquinone (28). A scheme depicting a possible pathway for RQ biosynthesis in *C. elegans* is shown in Fig. 5, which utilizes COQ-2, COQ-3, COQ-5, and COQ-6. The order of these proposed steps will need to be determined.

**Figure 5.**
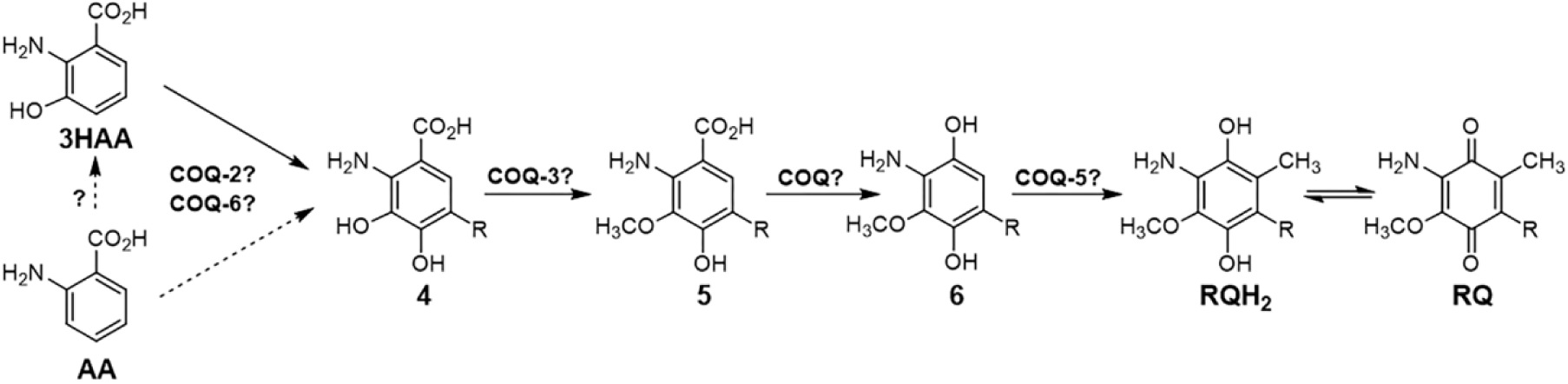
Proposed pathway for RQ biosynthesis in *C. elegans*. Either 3HAA or AA are proposed to be arylamine precursors to RQ. The Q biosynthetic enzymes, COQ-2 and COQ-6, may be used to form the common precursor, 2-amino-3,4-dihydroxy-5-nonaprenylbenzoic acid (compound 4). *O*-methylation of compound 4 can be achieved using a SAM-dependent methyltransferase, most likely COQ-3. The resulting compound 5 must be decarboxylated and hydroxylated, respectively, using the COQ enzyme(s), which would be analogous to those used in the Q biosynthetic pathway, to form the final 1,4-hydroquinone precursor to RQH_2_ (compound 6). The COQ-5 *C*-methyltransferase is proposed to catalyze the final methylation step to form RQH_2_, which can be oxidized to RQ. In *C. elegans*, “R” represents a tail with nine isoprenoid units (n=9).

Our results highlight the existence of two independent evolutionary pathways for RQ biosynthesis. Interestingly, *R. rubrum*, and protists that synthesize RQ, lack the kynurenine pathway and use Q and RquA for the biosynthesis of RQ. In contrast, KYNU-1 is present in all helminths and in bivalves, suggesting that the kynurenine pathway has been co-opted for RQ biosynthesis. Our findings have practical applications for the identification of potential targets in the RQ biosynthetic pathway for antihelminthic drug development. Parasitic helminth infections have become a global health epidemic, and in the face of emerging drug resistance, new treatments are necessary to combat them (29). A key issue regarding future studies is to understand why mammals, and other animals that possess the kynurenine pathway, do not synthesize RQ. The discovery of key enzymatic steps that discriminate between Q and RQ precursors will be highly relevant to target a metabolic pathway that is essential for helminth survival within the mammalian host under hypoxic conditions, such as those found in the intestine.

### Experimental Procedures

#### Caenorhabditis elegans *strains and culture conditions*

The *C. elegans* strains used in this study are listed in Table S2. Transgenic lines were obtained according to (30). The pRF4 plasmid containing the injection marker *rol-6(su1006)* was co-injected with constructs containing *Pkynu-1*::*kynu-1*::*gfp* cloned into the pPD95.77 plasmid, and injected into *kynu-1(tm4924)* animals. Independent transgenic lines were isolated and observed. The general methods used for culturing and maintenance of *C. elegans* are described in (31). All chemical reagents were purchased from Sigma-Aldrich (St. Louis, MO, USA). Chemical supplementation was carried out adding 10 mM AA or 10 mM 3HAA to NGM agar plates.

#### Reporter construct for expression and localization analysis

The expression pattern of *kynu-1* was determined using GFP as a reporter. The translational constructs *Pkynu-1::kynu-1*::*gfp* included promoter (1.4 kb), exons and introns of *kynu-1* in frame with the *gfp* coding sequence Sequences were amplified by PCR using appropriate primers (Table S3), from N2 genomic DNA. The PCR products were cloned into the pPD95.77 vector that provides the *unc-54* 3’UTR. For the study of the *kynu-1* expression pattern under hypoxic conditions, adult worms of the transgenic lines expressing the construct *Pkynu-1*::*kynu-1*::*gfp* were grown at 0.42% oxygen, 20 °C during 20 h in a C-Chamber incubator with a ProOx 110 oxymeter (Biospherix, Parish, NY, USA). Worms were immediately mounted for visualization under the microscope. Animals were visualized under a confocal microscope Zeiss LSM 880 and images captured with the Zen black 2.3 software and processed with Fiji (32). Embryos were obtained by a transverse cut in a gravid adult (early stages) or picked directly from the plate (late embryonic stages).

#### RNA interference assay

The expression of *C. elegans kynu-1, coq-3, coq-5, coq-6*, and *coq-7 (clk-1)* genes were interfered by the *E. coli* strain HT115 containing the plasmid pL4440 encoding the gene of interest (Table S4). Plasmids without an insert DNA (EV) or encoding *unc-22* were used as controls. *E. coli* strains were grown overnight at 37 °C in LB plus ampicillin (50 µg/mL) and carbenicillin (30 µg/mL), followed by a 2 h outgrowth to obtain a cell density of 0.4-0.6 OD_600_ units. Each strain was seeded onto 20 NGM agar plates (150 µL per plate) plus ampicillin, carbenicillin and 1 mM IPTG (to induce expression of dsRNA) and incubated for 48 h at 37 °C. RNAi was carried out by plating *C. elegans rrf-3*(pk1426), which were age synchronized to the L1 stage, onto the seeded *E. coli* plates at 22 °C and grown for 7 days. Worms were washed from plates with M9 buffer, divided into aliquots for pelleting, and frozen at −80 °C until use.

#### Lipid extraction

For lipid extractions of *C. elegans* N2 or mutant strains, 4,000 synchronized L1s were grown on NGM plates at 20 °C to adulthood. For each experiment, approximately 10,000 adult worms were harvested and washed several times with 18 MΩ water to obtain pellets for extraction (∼100 mg). Lipid extraction of *rrf-3* strains from RNAi assays were also performed using pellets containing ∼100 mg of worms, prepared from feeding plates as described above. Prior to extraction, 1000 pmol Q_3_ internal standard was added to pellets, and then lipids were extracted using hexanes and ethanol as previously described (12).

#### RNA isolation and RT-qPCR

RNA was extracted from ∼100 mg worm pellets using TRIzol reagent and the Zymo Quick-RNA MiniPrep Kit and further purified using the Zymo RNA Clean and Concentrator Kit (Zymo Research, Irvine, CA, USA). cDNA was prepared using the High Capacity RNA to cDNA kit (Applied Biosystems, Waltham, MA, USA) and TaqMan gene expression assays (Table S4) were optimized and performed for each RNAi strain as previously described (10) using the endogenous control assay, *cdc-42* (Ce02435138_g1).

#### LC-MS quantitation

LC-MS samples were prepared as described in (12). Standards were prepared and extracted at the following concentrations: Q_3_ (10 pmol/10 µL injection) and RQ_9_ (0.75, 1.5, 3.0, 4.5, or 6.0 pmol/10 µL injection). The RQ_9_ standard was isolated from *Ascaris suum* lipid extracts at Gonzaga University. In the absence of a standard, the quantity of Q_9_ was determined using a pmol conversion from the RQ_9_ standard curve and applying a RQ/Q response correction factor of 2.45 determined from RQ_10_/Q_10_ and RQ_8_/Q_8_ standard curves (12). Additional quinone-specific parameters are listed in Table S5. Samples were analyzed in triplicate and the pmol quinone was determined from the standard curve and corrected for recovery of internal standard. Samples were normalized by mg pellet mass.

## Acknowledgements

The authors would like to thank Hugo Bisio, Cecilia Martínez, Gastón Risi, Drs. Lucía Otero and Jorge Pórfido (Laboratorio de Biología de Gusanos) for helpful discussions on RQ biosynthesis, as well as Drs. Jeff Cronk and Kirk Anders from Gonzaga University (GU), Dr. Catherine Clarke (UCLA), Drs. Andrew Roger, Courtney Stairs and David Langelaan (Dalhousie University), NS and Dr. Gilles Basset and Ann Bernert (University of Florida, Gainesville). We thank Dr. Jennifer Watts (WSU) for RNAi clones and Alex Poppel for his high school summer volunteer work at GU.

## Abbreviations

ETC: electron transport chain
HA: anthranilic acid
3HAA: 3-hydroxyanthranilic acid
HKYN: 3-hydroxykynurenine
KYN: kynurenine
Q: ubiquinone
RQ: rhodoquinone.

## Supporting Information

### List of Materials

p. S-2, TABLE S1. Statistical analysis of RQ_9_ and Q_9_ levels in mutant strains and RNAi knockdowns

p. S-3, TABLE S2. *C. elegans* strains used in this study

p. S-4, TABLE S3. Primers used for *kynu-1* reporter construction

p. S-5, TABLE S4. RNAi clones and TaqMan assays for RT-PCR

p. S-6, TABLE S5. LC-MS parameters for each quinone

p. S-7, **Figure S1.** *kynu-1* is expressed during embryogenesis and the first larval stage in hypodermis and intestinal cells

p. S-8, **Figure S2.** Quantitation of gene expression from *C. elegans* RNAi strains using RT-qPCR

**TABLE S1.** Statistical analysis of RQ_9_ and Q_9_ levels in mutant strains and RNAi knockdowns

| Strain | Avg pmol<br>RQ<sub>9</sub>/mg pellet | p value<sup>a</sup><br>(N = 3) | Avg pmol<br>Q<sub>9</sub>/mg pellet | p value<br>(N = 3) |
| --- | --- | --- | --- | --- |
| N2 | 3.27 ± 0.15 |  | 17.03 ± 0.71 |  |
| *afmd-1* | 1.51 ± 0.09 | < 0.001 | 13.24 ± 1.80 | 0.0138 |
| *kynu-1* | 0 | < 0.001 | 19.27 ± 1.76 | 0.0633 |
| *kmo-1* | 0.51 ± 0.07 | < 0.001 | 13.49 ± 0.88 | 0.0027 |
| *haao-1* | 3.41 ± 0.61 | 0.360 | 14.41 ± 0.80 | 0.0065 |
| EV | 2.36 ± 0.20 |  | 9.48 ± 1.43 |  |
| *kynu-1* (RNAi) | 0.51 ± 0.05 | < 0.001 | 9.95 ± 2.52 | 0.397 |
| *coq-3* (RNAi) | 1.67 ± 0.12 | 0.003 | 7.01 ± 0.58 | 0.025 |
| *coq-5* (RNAi) | 1.26 ± 0.17 | < 0.001 | 5.21 ± 1.05 | 0.007 |
| *coq-6* (RNAi) | 1.02 ± 0.04 | < 0.001 | 5.10 ± 0.22 | 0.003 |
| *coq-7* (RNAi) | 2.05 ± 0.20 | 0.068 | 6.29 ± 0.32 | 0.010 |
| *unc-22* (RNAi) | 2.59 ± 0.60 | 0.283 | 9.48 ± 1.92 | 0.500 |

<sup>a</sup>The Student's T-test was used to analyze triplicate samples with significance noted at the $\alpha < 0.05$ level. The standard deviation in each data set is represented by ± and shown as error bars in Figs. 2B, 2C and 4B.

**TABLE S2.** *C. elegans* strains used in this study.

| Strain | Gene | Allele | Variation type | Nucleotide change | Genotype | Source |
| --- | --- | --- | --- | --- | --- | --- |
| N2 | *Bristol wild isolation* |  |  |  |  | CGC |
| NL2099 | *rrf-3* | pk1426 | deletion | 3055 bp deletion | *rrf-3(pk1426) II* | CGC |
| Tm4924 | *kynu-1* | tm4924 | insertion/deletion | 19 bp insertion<br>521 bp deletion | *kynu-1(tm4924) X* | NBPJ |
| Tm4529 | *kmo-1* | tm4529 | deletion | 326 bp deletion | *kmo-1(tm4529) V* | NBPJ |
| Tm4547 | *afmd-1* | tm4547 | deletion | 425 bp deletion | *afmd-1(tm4547) IV* | NBPJ |
| Tm4627 | *haao-1* | tm4627 | insertion/deletion | 9 bp insertion<br>305 bp deletion | *haao-1(tm4627) V* | NBPJ |
| IH25 |  |  |  |  | *kynu-1(tm4924)X*;<br>*Ex[Pkynu-1::kynu-1::gfp, pRF4]* | This study |

**TABLE S3.**
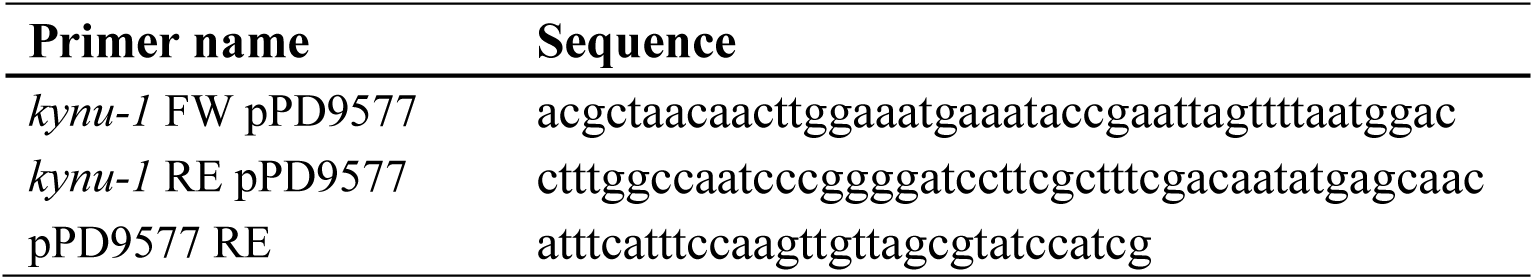
Primers used for *kynu-1* reporter construction

**TABLE S4.** RNAi clones and TaqMan assays for RT-PCR

| Strain | Gene | <sup>a</sup>Clone Number | Insert | Source | <sup>b</sup>TaqMan assay |
| --- | --- | --- | --- | --- | --- |
| *E. coli*<br>HT115<br>(DE3) | *kynu-1* | DFCIp3320G0510040D | C15H9.7 | Source Bioscience | Ce02495988_g1 |
|  | *coq3* | CUUkp3303J037Q | sjj_Y57G11C.11 | Source Bioscience | Ce02467843_g1 |
|  | *coq5* | CUUkp3302K054Q | sjj_ZK652.9 | Source Bioscience | Ce02449325_g1 |
|  | *coq6* | CUUKp3315A0214Q | sjj2_K07B1.2 | Source Bioscience | Ce02479593_g1 |
|  | *coq7/clk-1* | CUUkp3302B242Q | sjj_ZC395.2 | <sup>c</sup>gift | Ce02446729_g1 |
|  | *unc-22* | CUUkp3303K066Q | sjj_ZK617.1 | <sup>c</sup>gift | Ce02465425_g1 |

<sup>a</sup>All clones were from Ahringer library, L4440 (pPD129.36) except for *kynu-1* was from Vidal library, pL4440\_DEST
<sup>b</sup>Purchased from ThermoFisher Scientific (Rockford, IL, USA) with FAM-MGB, 20X
<sup>c</sup>Gift from Dr. Jennifer Watts, School of Molecular Sciences, Washington State University, Pullman, WA

**TABLE S5.** LC-MS parameters for each quinone

| **MS parameter** | **Q<sub>3</sub>** | **RQ<sub>9</sub>** | **Q<sub>9</sub>** |
| --- | --- | --- | --- |
| Dwell time (s) | 0.1 | 0.1 | 0.1 |
| Cone (V) | 20 | 39 | 35 |
| Collision (V) | 20 | 30 | 30 |
| Precursor mass [M+H]<sup>+</sup> (*m/z*) | 387.2 | 780.6 | 795.6 |
| Ion product mass [M]<sup>+</sup> (*m/z*) | 197.2 | 182.2 | 197.2 |

**Figure S1.**
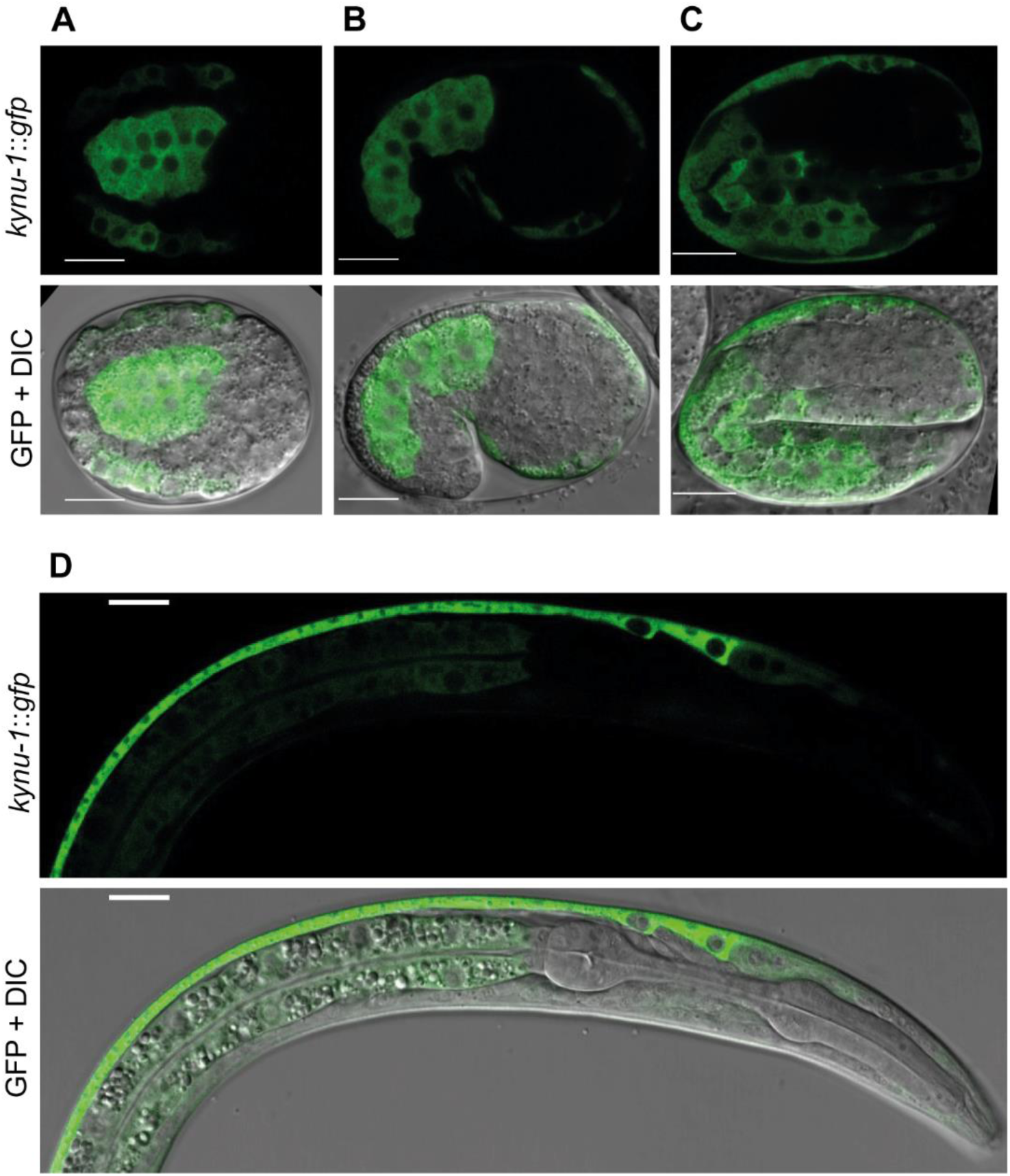
*kynu-1* is expressed during embryogenesis and the first larval stage in hypodermis and intestinal cells. Confocal images of selected planes show transgenic animals expressing the translational construct *Pkynu-1*::*kynu-1*::*gfp*. The stages shown are: (A) E16 dorsal view, (B) Comma lateral view, (C) 2-fold lateral view and (D) L1 lateral view. Scale bar 10 µm.

**Figure S2.**
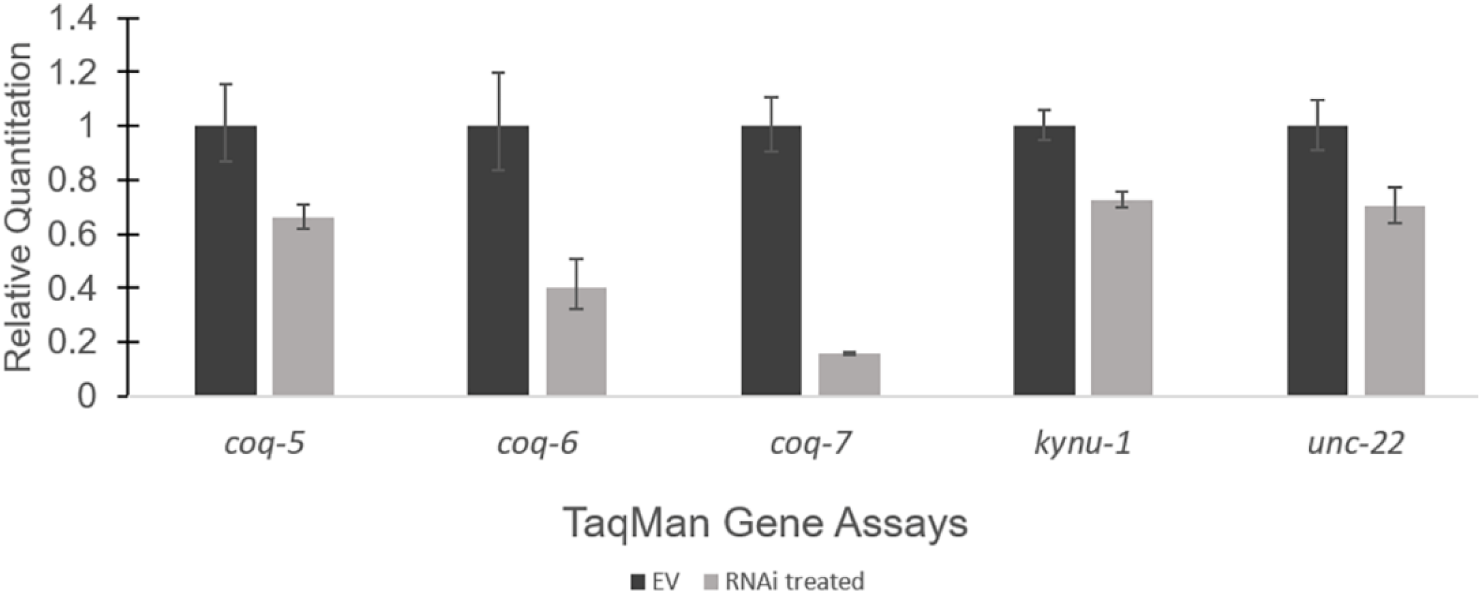
Quantitation of gene expression from *C. elegans* RNAi strains using RT-qPCR. A standard comparative C_T_ (ΔΔC_T_) experiment was performed with the TaqMan gene expression assays (Table S4) on a StepOnePlus™ Real-Time PCR system (Life Technologies, Waltham, MA). ROX™ dye was used as a passive reference and EV cDNA was used as an active reference in each experiment. Each cDNA sample and no RT control was tested in triplicate with each assay, and the average C_T_ values were generated for each biological sample with each gene target. ΔC_T_ and ΔΔC_T_ values were obtained in order to determine the range of fold-change values, comparing RNAi knockdowns to EV control. Relative quantitation (RQ) ranges were determined through standard propagation of error. TaqMan gene assays for *coq-5, coq-6, coq-7, kynu-1* and *unc-22* showed reduction of expression compared to the EV reference, using the *cdc-42* endogenous control. The relative quantitation values were significant for *coq-6* and *coq-7* (more than 2-fold smaller), and weakly significant for *coq-5, kynu-1*, and *unc-22*. The *coq-3* TaqMan assay did not allow for quantitation due to inconsistent amplification of *cdc-42* in the sample and reference.

## FOOTNOTES

This study was supported by the Agencia Nacional para la Innovación y la Investigación ANII FCE_1_2014_104366 (GS), FOCEM (MERCOSUR Structural Convergence Fund) COF 03/11 (Institut Pasteur de Montevideo), Howard Hughes Medical Institute through the Undergraduate Science Education Program (Gonzaga University), the Dr. Scholl Foundation (JNS), and the Agencia Nacional de Innovación e Investigación post-doctoral fellowship PD_NAC_2013_11008 (IC). The Caenorhabditis Genetics Center is funded by NIH Office of Research Infrastructure Program: P40 OD010440.

